# Pep-PU-GAN: Positive-Unlabeled Adversarial Learning for Peptide Function Prediction

**DOI:** 10.64898/2026.09.13.751209

**Authors:** Farzad Midjani, Samaneh Hashemi, Fateme Zahra Keshtkar, Mahdi Malekpour, Bahar Saberzadeh Ardestani, Bardia Khosravi

**Affiliations:** Systems Medicine Research Core, Shiraz University of Medical Sciences, Shiraz, Iran; Student Research Committee, Shiraz University of Medical Sciences, Shiraz, Iran; Department of Medical Biotechnology, School of Advanced Medical Sciences and Technologies, Shiraz University of Medical Sciences, Shiraz, Iran; Department of Internal Medicine, Yale School of Medicine, New Haven, CT, USA; Department of Radiology, Mayo Clinic, MN, USA; Department of Radiology, Yale University, New Haven, CT, USA

**Keywords:** Positive-unlabeled learning, Generative adversarial networks, Graph neural networks, Peptide classification, Neuropeptide prediction, Self-training

## Abstract

Peptide classification remains challenging in bioinformatics because of limited labeled data, particularly the scarcity of verified negative examples, and the complex relationship between amino acid sequences and biological functions. This study introduces Pep-PU-GAN, a deep learning framework that combines positive-unlabeled (PU) learning, generative adversarial networks (GANs), and graph neural networks (GNNs) for peptide classification. Peptides are represented as sequence-derived residue graphs, with amino acids as nodes and edges connecting adjacent residues, enabling attention-based message passing over local neighborhoods. The architecture includes a generator that produces synthetic peptide embeddings in encoder space and a dual-function discriminator that distinguishes real from synthetic embeddings while performing PU classification. Training uses a custom loss integrating non-negative PU (nnPU) risk estimation with adversarial objectives. A self-training mechanism further incorporates high-confidence synthetic positive embeddings to augment the training set and improve performance. Evaluated on neuropeptide classification using 4,049 positive neuropeptides and 8,558 unlabeled peptides, Pep-PU-GAN outperformed baseline models, achieving an F1 score of 0.93 and an AUROC of 0.98 on an independent held-out benchmark. Pep-PU-GAN provides a promising approach for peptide classification tasks with scarce labeled and abundant unlabeled data, with potential applications in computational biology and drug discovery.

## 1 Background

Peptides are short amino-acid chains involved in diverse biological processes, including cellular signaling, host defense, regulatory activity, and intercellular communication [1]. Among these molecules, neuropeptides represent an important functional class that acts in the nervous system as neurotransmitters, neuromodulators, and signaling molecules [2]. Reliable peptide identification can support the study of peptide-mediated signaling and help prioritize candidates for functional or therapeutic investigation. However, experimental peptide annotation is labor-intensive, time-consuming, and difficult to scale. As peptide databases continue to expand, computational prediction has become an important tool for prioritizing candidate peptides for further biological investigation [3, 4].

Deep-learning models have achieved strong performance in peptide-sequence classification [5]; however, most supervised approaches require labeled examples from both the positive and negative classes. This requirement is poorly suited to peptide-function prediction because experimentally confirmed positives are more readily available than verified negatives, and unannotated peptides cannot be assumed to lack the target function. Treating such sequences as negatives introduces label noise and benchmark bias, particularly when annotation coverage differs across species, peptide families, databases, and experimental settings [6]. Furthermore, known positives often represent only a limited portion of the functional sequence space, whereas the unlabeled pool contains a mixture of irrelevant sequences and undiscovered positives [7, 8]. An appropriate learning framework must therefore extract information from the unlabeled pool without assigning all unannotated peptides to the negative class.

Positive-unlabeled (PU) learning addresses peptide classification problems in which experimentally confirmed positives are available but reliable negative labels are scarce. Instead of assigning negative labels to all unannotated sequences, PU learning estimates classification risk from labeled positives and an unlabeled mixture that may contain both positive and negative instances [9, 10]. Building on this principle, we introduce Pep-PU-GAN and apply it to neuropeptide identification. The principal contribution is an integrated framework comprising: (i) non-negative PU risk minimization for training without curated negative labels; (ii) a residue-adjacency graph encoder coupled to a dual-branch discriminator and latent-space adversarial augmentation [11, 12, 13]; and (iii) a generator-assisted self-training procedure that incorporates confidence-filtered synthetic latent vectors as pseudo-positive signals during later training stages [14].

## 2 Methods

The overall workflow of the proposed Pep-PU-GAN framework is illustrated in Fig. 1.

**Fig. 1.**
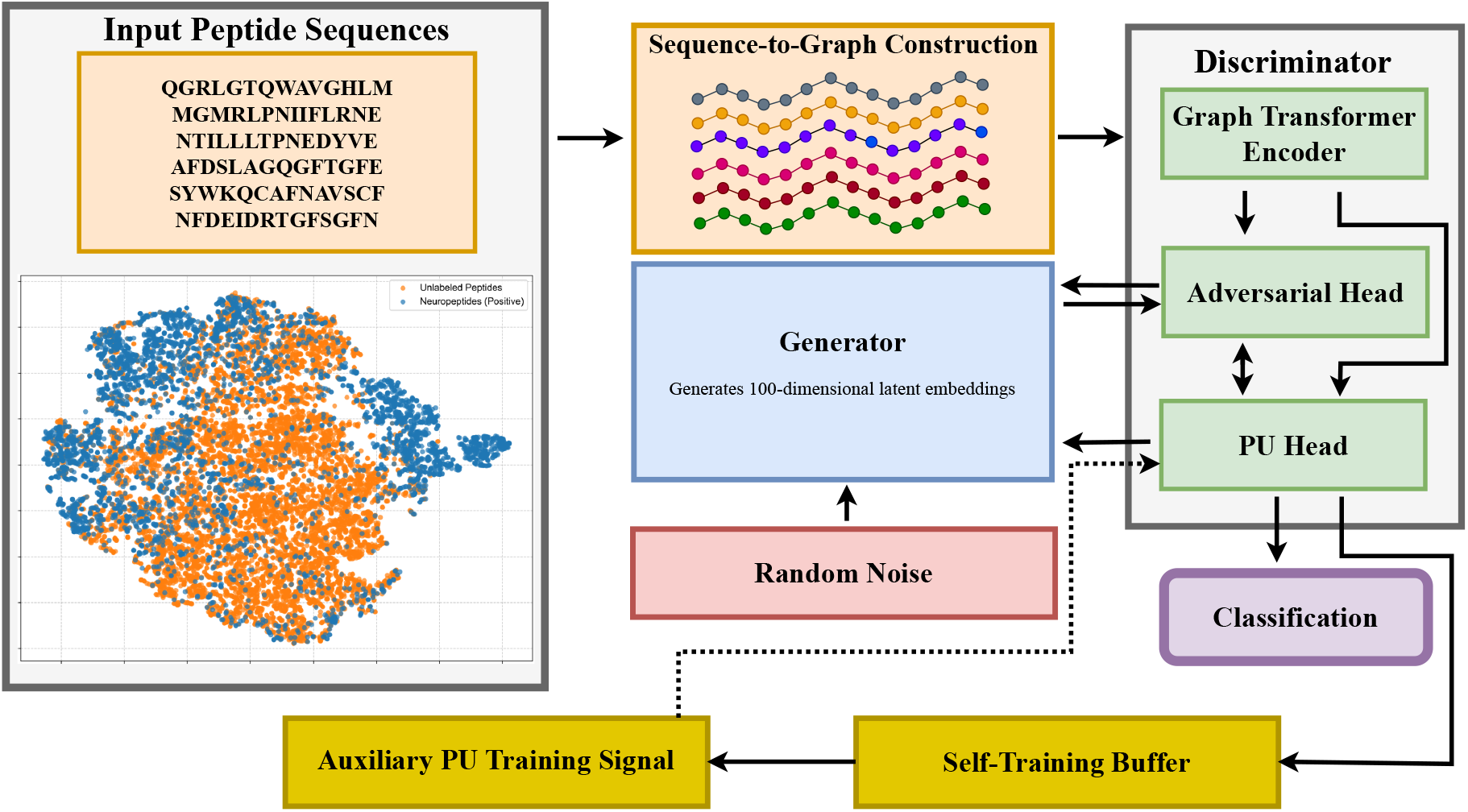
Overview of the Pep-PU-GAN workflow. The embedded projection visualizes the distribution of labeled neuropeptides and unlabeled peptides based on sequence-derived peptide features. Peptide sequences are converted into sequence-derived residue graphs, encoded by a graph transformer, and processed by a dual-branch discriminator for adversarial discrimination and PU classification. The generator maps random noise to 100-dimensional latent embeddings, which support adversarial training and late-stage self-training through an auxiliary PU training signal.

### 2.1 Data Collection Methodology

We compiled positive and unlabeled datasets to evaluate Pep-PU-GAN on neuropeptide identification. For the positive dataset, experimentally validated neuropeptides were obtained from NeuroPep 2.0 [15]. The initial corpus from NeuroPep 2.0 comprised 11,417 sequences. CD-HIT software reduced sequence redundancy [16] with a similarity threshold of 0.9, ensuring a non-redundant set of positive examples. The unlabeled dataset used for PU training consisted of 8,558 filtered Cow Peptides sourced from the PeptideAtlas database [17].

For external benchmark construction, a separate *C. elegans* peptide pool containing 106,414 sequences was obtained from PeptideAtlas as the source of proxy-negative benchmark candidates. After removing exact sequence overlaps with the positive and PU-training unlabeled sets, 106,189 external benchmark candidates remained.

Both the positive and training-unlabeled peptide sets underwent a final filtering step. Peptides were retained if they consisted exclusively of the 20 standard (canonical) amino acids and had lengths of 6–50 amino acids. This final filtering resulted in a Positive set (P) of 4,049 neuropeptide sequences and an Unlabeled set (U) of 8,558 peptide sequences. The external *C. elegans* pool was processed using the same sequence-validity criteria, but it was never used as PU training data or for model fitting.

### 2.2 Experimental Setup

#### 2.2.1 Data Partitioning

Labeled positives were divided into an outer training set and an outer held-out test set. The training positives were further divided into a cross-validation pool and a reserved Fold 6 calibration subset. The external benchmark candidate pool was split once into validation/calibration and held-out test candidate pools. The PU-training unlabeled set was used only during model fitting and was never treated as the negative class during evaluation.

#### 2.2.2 External Proxy-Negative Benchmark Construction

From the external *C. elegans* candidate pool, proxy negatives were selected separately for each validation, calibration, and final held-out benchmark. Negative selection used only the training-side positives available for the corresponding split, so target positives and final held-out candidates did not contribute to the proxy-negative selection rule.

Candidate peptides were first screened for sequence dissimilarity to the reference positive set using character 2- and 3-mer TF-IDF representations and the three nearest reference-positive neighbors. Candidates within a gray zone of 0.10 around the similarity threshold were further evaluated using normalized Levenshtein similarity. Candidates were retained only when their similarity to the reference positives remained below a threshold of 0.40. The surviving candidates were then scored by distance from a positive-only manifold fitted on the reference positives using peptide length, amino acid composition, and dipeptide composition features. PCA retained 95% of the variance with a maximum of 32 components, followed by Ledoit–Wolf Mahalanobis-distance estimation. Distance cutoffs were evaluated sequentially at positive-distance quantiles of 0.99, 0.975, 0.95, and 0.90.

A reliability score was computed by combining the manifold-distance rank and sequence-dissimilarity rank with weights of 0.60 and 0.40, respectively. If the distance-quantile filters yielded insufficient candidates, a global reliability ranking was used. Final proxy negatives were selected deterministically to match the length profile of the target positive split at a 1:1 proxy-negative-to-positive ratio. For Folds 1–5, fold-training positives defined the proxy-negative selection rule and fold-validation positives defined the target benchmark. Fold 6 used all cross-validation positives as the reference set and the reserved calibration positives as the target benchmark. The final held-out benchmark used all training-side positives as the reference set and the outer held-out positives as the target benchmark. Final held-out proxy-negative candidates were separated from validation, calibration, model-selection, and threshold-tuning steps to limit evaluation leakage.

#### 2.2.3 Evaluation Protocol

Model evaluation comprised five-fold cross-validation (Folds 1–5) and a separate calibration run designated as Fold 6. In Folds 1–5, the cross-validation positive pool was partitioned into fold-specific training and validation positives. Fold 6 trained on the complete cross-validation positive pool and was validated on the reserved calibration positives. In every fold, training used the corresponding positives together with the full unlabeled PU-training set, and validation paired the fold positives with a balanced set of proxy negatives mined from the validation-benchmark candidate pool. For each fold, the decision threshold was selected from the corresponding validation benchmark. The final operating threshold was then fixed as the median of the six validation-selected thresholds. The final held-out test benchmark was constructed only once by pairing the outer held-out positive peptides with proxy negatives mined from the reserved held-out test-benchmark candidate pool, and final test performance was reported using an ensemble of all six trained models obtained by averaging predicted probabilities.

#### 2.2.4 Evaluation Metrics

The performance was evaluated using standard binary classification metrics: Precision, Recall, F1 score, AUROC, and Accuracy. For reporting these metrics, validation and test performance were computed on balanced benchmarks composed of held-out positives and proxy negatives selected from the external peptide pool.

For all final held-out evaluations, 95% confidence intervals for threshold-dependent metrics were estimated using 10,000 stratified bootstrap resamples, with validation-derived thresholds kept fixed.

### 2.3 Graph Representation of Peptides

Each peptide sequence in both the positive and unlabeled datasets is converted into a sequence-derived residue graph. Amino acids correspond to nodes, and a node stores an integer token for the residue type (20 canonical amino acids), which is mapped to a learnable embedding. Bidirected edges are added only between adjacent residues in the primary sequence, preserving local sequential adjacency for message passing [18, 19]. This graph construction provides a residue-level relational representation in which an amino acid is encoded in the context of its immediate neighbors. After graph convolution and pooling, the resulting peptide embedding therefore reflects local sequence organization and neighborhood-dependent residue effects rather than amino-acid identity alone.

### 2.4 Model Architecture: Pep-PU-GAN

Pep-PU-GAN comprises a shared peptide encoder, a latent-space generator, and a discriminator with adversarial and PU-classification branches.

#### 2.4.1 Graph Transformer Encoder

The graph transformer encoder (Fig. 2) is a shared feature extractor that converts peptide graphs into 100-dimensional latent embeddings. It starts with an amino acid embedding layer mapping node indices to 50-dimensional vectors, followed by a single TransformerConv layer from PyTorch Geometric with one attention head, a 100-dimensional output, and dropout of 0.3 for context-dependent message passing [20, 21, 22]. The node features are then processed by layer normalization, Leaky ReLU activation (slope 0.2), and dropout (0.3). Global mean pooling yields a 100-dimensional graph-level representation, which is finalized with batch normalization for batch-wise consistency [23, 24, 25].

**Fig. 2.**
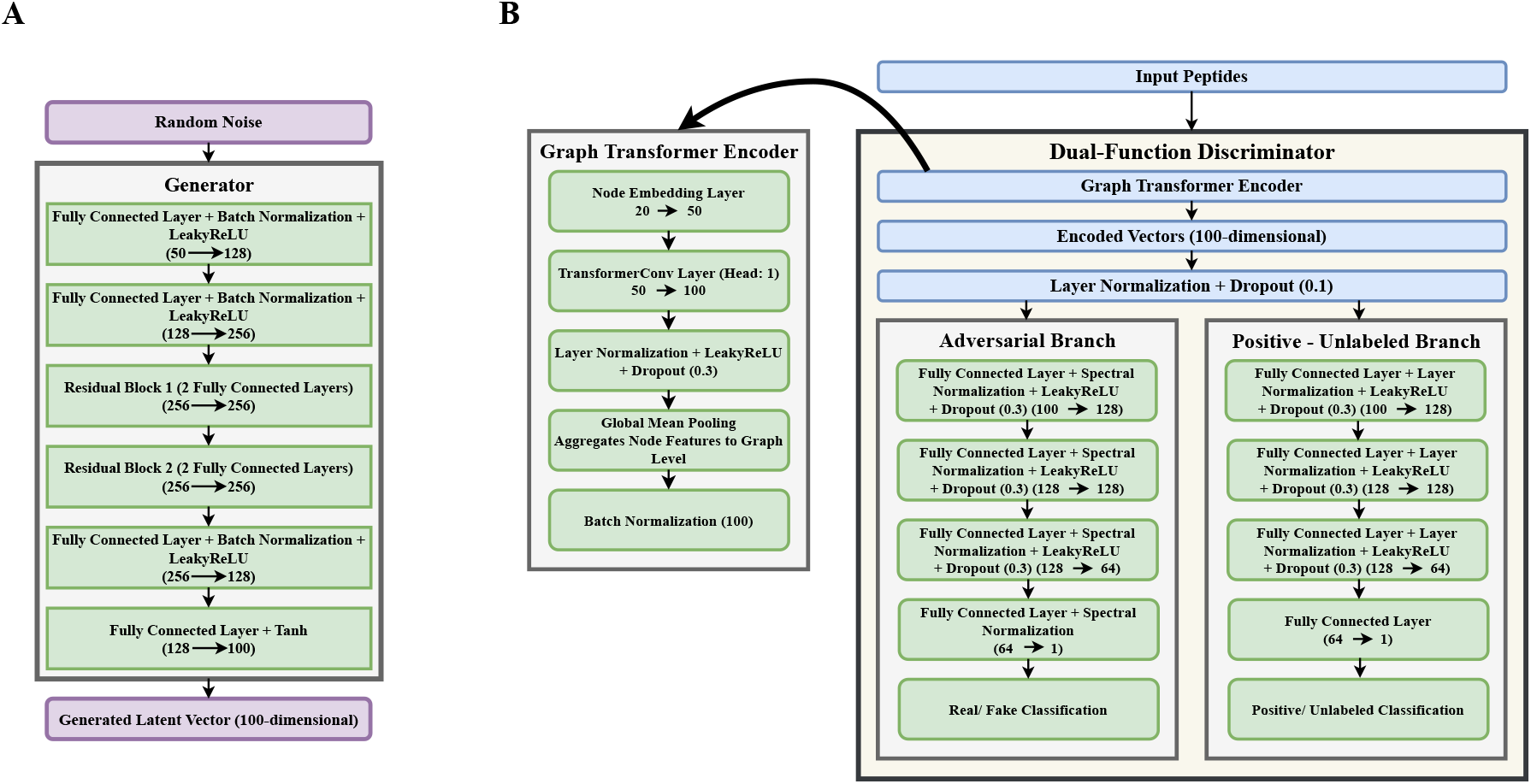
Architecture of the generator and dual-function discriminator in Pep-PU-GAN. (A) Generator architecture, mapping 50-dimensional random noise to 100-dimensional latent vectors. (B) Dual-Function Discriminator architecture, processing 100-dimensional latent vectors through latent normalization and two task-specific branches for adversarial discrimination and PU classification.

#### 2.4.2 Generator

As shown in Fig. 2, the generator aims to learn the distribution of real peptide latent embeddings, i.e., the 100-dimensional vectors produced by the graph transformer encoder. It is designed to map random noise vectors, sampled from a 50-dimensional standard normal distribution, to synthetic latent vectors that mimic these real embeddings. Architecturally, it is a residual multilayer perceptron: two fully connected blocks with batch normalization and Leaky ReLU are followed by two 256-dimensional residual blocks, a projection back to 128 dimensions, and a final linear layer with Tanh activation. The generator therefore outputs continuous 100-dimensional latent vectors that are passed directly to the discriminator.

#### 2.4.3 Dual-Function Discriminator

As illustrated in Fig. 2, the discriminator processes both real peptide graphs and synthetic latent vectors through two task-specific branches. Real peptide graphs are first encoded by the graph transformer encoder, whereas generator-produced latent vectors bypass this encoding step. In both cases, the resulting 100-dimensional latent representation is processed by latent layer normalization and input dropout (0.1) before entering the downstream heads.

##### Adversarial Branch

The adversarial branch distinguishes encoder-derived peptide embeddings from generator-produced latent embeddings. It uses a fully connected architecture, with Leaky ReLU activation (slope 0.2) and dropout (0.3) after each hidden layer. Spectral normalization is applied to each linear layer in the adversarial branch to control its operator norm during adversarial optimization [26]. The 64-dimensional hidden representation immediately preceding the final logit is also used for feature-matching loss computation.

##### PU Classification Branch

As shown in Fig. 2, the PU branch receives the same 100-dimensional latent vector as the adversarial branch and outputs a single logit for positive-class prediction. Each hidden layer is followed by layer normalization, Leaky ReLU activation (slope 0.2), and dropout (0.3). The final bias of the PU head is initialized from a fold-specific estimate of the positive-class prior to calibrate early-training probabilities.

This dual-branch architecture allows the adversarial and PU objectives to share the graph encoder while maintaining task-specific downstream representations. As a result, the generator is encouraged not only to produce realistic latent embeddings for the adversarial branch, but also to generate samples that receive high positive confidence from the PU branch. In the later stage of training, a curriculum-controlled attenuation factor is applied only to the adversarial path for generated latents, reducing excessive interference between adversarial updates and PU classification.

### 2.5 Self-Training with Generator-Produced Samples

After the PU warm-up and GAN-integration stages, Pep-PU-GAN applies a confidence- and diversity-filtered self-training procedure to generated latent vectors:

1. After the initial PU-only warm-up and a short GAN stabilization period, the generator produces batches of synthetic latent vectors from random noise.
2. These generated latent vectors are passed through the PU classification branch of the discriminator, which assigns each sample a positive-class confidence score [27, 28].
3. Generated samples are retained for self-training only if they satisfy a confidence threshold of at least 0.75. To avoid degenerate reuse of highly similar samples, a diversity constraint is enforced: a new latent vector is added only if its *L*_2_ distance from the existing buffer contents exceeds 0.15. In addition, once sufficient buffer statistics are available, generated samples are required to remain within the empirical distribution of the stored latent features. The self-training buffer stores up to 5,000 latent vector-confidence pairs.
4. During later training epochs, when the self-training buffer is nonempty, generated latent vectors that have passed the confidence, diversity, and distribution filters are sampled from *B*_t_ and assigned pseudo-positive labels. The real peptide graphs and generated latent vectors are processed separately by the PU branch, after which their PU logits and labels are concatenated. The nnPU objective is then evaluated on this combined set, causing the generated samples to contribute to the empirical positive terms of the PU risk. The resulting loss is multiplied by the standard PU-loss weight and used in an additional discriminator optimization step before the regular generator and discriminator updates.

### 2.6 Training Objectives

As illustrated in Fig. 3, Pep-PU-GAN jointly optimizes generator and discriminator objectives that combine adversarial training with nnPU risk for PU classification [10].

**Fig. 3.**
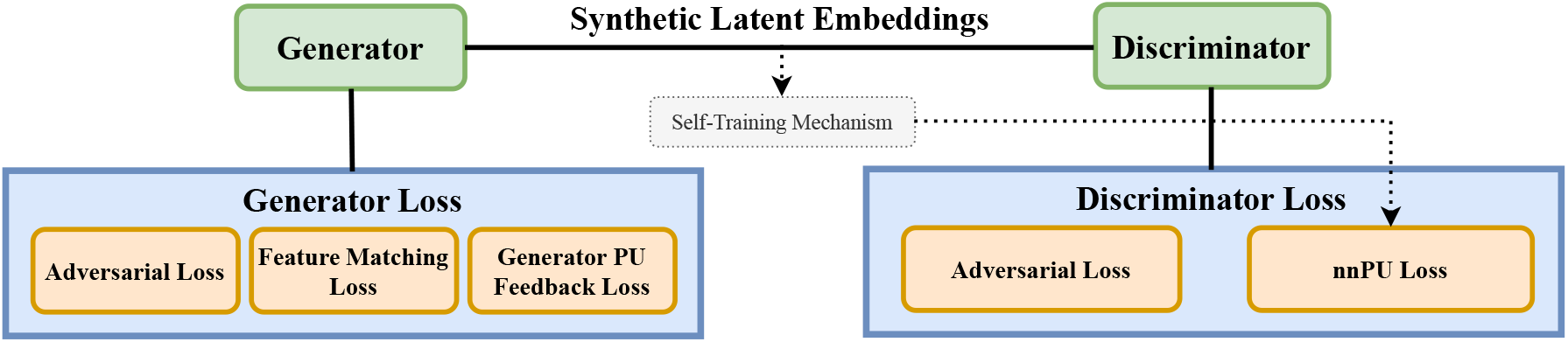
Training objectives used in Pep-PU-GAN. The discriminator combines the nnPU classification objective with an adversarial hinge objective, whereas the generator combines adversarial, feature-matching, and PU-feedback terms.

#### 2.6.1 Discriminator Loss

The discriminator’s total loss in Pep-PU-GAN combines an adversarial loss and an nnPU loss to balance generative and classification objectives. The adversarial loss, applied to the discriminator’s adversarial branch, uses the hinge formulation to distinguish real peptide embeddings from generator-produced latent vectors. The PU loss targets the PU classification branch and follows an adapted nnPU risk decomposition with non-negative correction [10]. During optimization, its positive-risk terms are scaled by an adaptive weight *w*_t_, initialized from the training-only class-prior estimate.

Let *z* denote a real latent embedding and 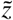 a generated latent embedding. Let *D*_adv_ (·) denote the adversarial logit and *D*_pu_ (·) the PU-branch logit. The discriminator hinge adversarial loss is:

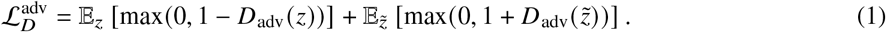

Following the nnPU formulation [10], we use the logistic surrogate losses

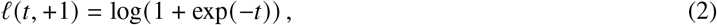

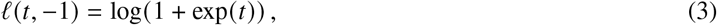

and define, at training epoch t,

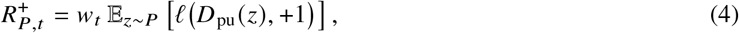

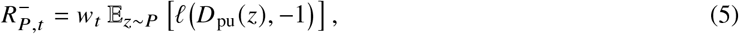

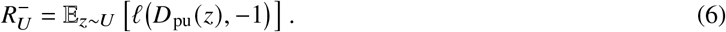

The resulting validation-guided nnPU objective is

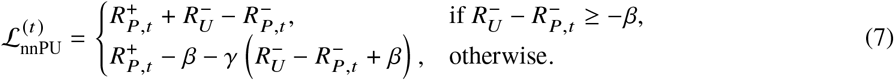

where *β* and *γ* are the nnPU correction parameters; in all experiments, *β* = 0 and *γ* = 1. The total discriminator objective at training epoch t is

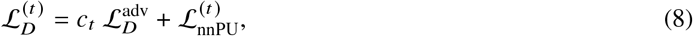

where *c*_t_ is the curriculum coefficient: *c*_t_ = 0 during the initial PU-only warm-up, increases linearly during GAN integration, and reaches 1 in the final stage of training.

#### 2.6.2 Generator Loss

The generator objective contains three terms: adversarial loss, feature-matching loss, and PU feedback loss. The adversarial term encourages generated embeddings to receive high real-sample scores from the discriminator. The feature-matching term reduces differences between the mean and variance of penultimate adversarial-layer activations for real and generated batches, following the stabilization rationale of feature matching [29]. The PU feedback term encourages generated embeddings to receive high positive-class scores from the PU branch. The total generator loss is the weighted sum of these terms.

Concretely, the generator hinge adversarial loss is:

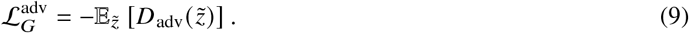

Let *f* (·) denote the discriminator feature vector used for feature matching. Matching both mean and variance yields:

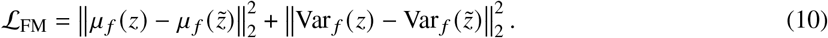

A PU feedback term that encourages 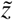 to be classified as positive is:

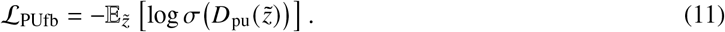

The overall generator loss is

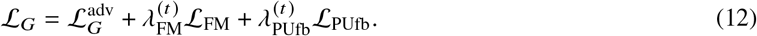

In all experiments, the base weights were set to *λ*_FM_ = 1.0 and *λ*_PUfb_ = 0.5.

### 2.7 Training and Implementation Details

The Pep-PU-GAN model was trained using separate Adam optimizers for the generator and discriminator [30], with learning rates of 1 × 10^*−*4^ and 2 × 10^*−*4^, respectively, following the two-time-scale update rationale for GAN training [31]. Both optimizers used a weight decay of 1 × 10^*−*5^ and beta parameters of 0.5 and 0.999. Cosine annealing learning-rate schedulers with warm restarts were applied to both optimizers [32]. To reduce the risk of exploding gradients, norm-based gradient clipping with a maximum norm of 1.0 was applied to both generator and discriminator parameters during backpropagation [33].

A curriculum learning strategy is implemented to manage the complexity of training the combined PU and GAN framework by gradually introducing the adversarial objectives. During the first 8 epochs, training is GAN-free: the encoder and PU branch are optimized using nnPU loss on real positive and unlabeled data, whereas the generator and adversarial branch remain inactive. During the subsequent 15 epochs (epochs 9–23), GAN training is introduced progressively. Specifically, the discriminator adversarial loss is multiplied by a linearly increasing curriculum coefficient, while the generator-side auxiliary terms, i.e., feature matching and PU feedback, are scaled from zero to their full weights. Self-training begins after the stabilization period has started; in our experiments, high-confidence generated latent embeddings were added to the buffer from epoch 14 onward. After the curriculum phase, the full GAN-PU objective is used. In this later stage, an attenuation factor on the adversarial path for generated latents is gradually increased by 0.05 per epoch up to 0.5, reducing excessive interference between adversarial updates and PU classification while leaving the PU branch itself unchanged.

For each training split, the initial positive-class prior for the nnPU loss is estimated exclusively from the training-side positive and unlabeled peptides using the Elkan–Noto label-frequency approach with out-of-fold logistic-regression predictions on handcrafted sequence features (peptide length, amino acid composition, and dipeptide composition) [7]. Let *S ∈* {0, 1} denote the observed PU label, where *S* = 1 for labeled positives and *S* = 0 for unlabeled samples. With training-side positive set *P* and unlabeled set *U*, the observed label frequency is

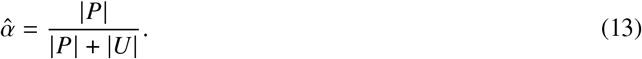

A logistic-regression selector *q* (*x*) ≈ *P S* = 1 *x* is fitted using *K*-fold out-of-fold predictions, and the labeling probability for positives is estimated as

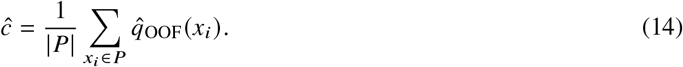

The training-only initial positive-class prior is estimated as

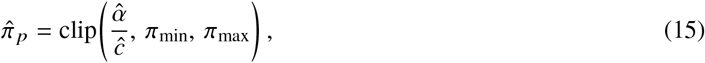

where *π*_min_ = 0.01 and *π*_max_ = 0.95. This estimate was used to initialize both the PU-head bias and the adaptive positive-risk weight, 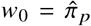; during training, the adaptive weight was constrained to 0.005, 0.95 . During optimization, *w*_t_ was updated from the precision–recall balance on the proxy-validation benchmark. When the precision-to-recall ratio exceeded 1.2, *w*_t_ was increased by 5% following an improvement in validation F1 and by 10% otherwise. When this ratio fell below 0.8, *w*_t_ was decreased by 5% following an improvement in validation F1 and by 10% otherwise. No update was applied when the ratio remained between 0.8 and 1.2, and *w*_t_ was constrained to the interval 0.005, 0.95 . Thus, *w*_t_ is a validation-guided risk coefficient and is not interpreted as an updated estimate of positive prevalence in the unlabeled set. Model selection is performed on balanced validation benchmarks composed of validation positives and proxy negatives, and the PU head is evaluated on these benchmarks using binary cross-entropy loss on logits. Early stopping prevents overfitting by halting training after 10 epochs without validation F1 improvement, and the best-F1 checkpoint is used for downstream evaluation [34]. Threshold selection is carried out strictly on validation/calibration benchmarks. The Pep-PU-GAN framework was implemented using PyTorch version 2.5.0 [35] and PyTorch Geometric version 2.6.1 [20].

### 2.8 Ablation Models

We evaluated two matched ablation baselines to quantify the performance differences associated with the adversarial/self-training module group and the graph encoder. The first was a GAN-free PU baseline designed to assess the effect of adversarial learning and generator-assisted self-training. This model retained the graph transformer encoder and PU classification branch of Pep-PU-GAN but removed the generator, adversarial branch, adversarial losses, feature-matching loss, PU feedback loss, and self-training with generated latent samples. All other training and evaluation settings, including the validation-guided nnPU objective, class-prior initialization, positive-risk-weight adaptation rule, validation procedure, and threshold-selection protocol, were kept identical to the full model.

The second was a graph-free sequence baseline designed to assess the contribution of sequence-derived graph modeling. In this model, the graph transformer encoder was replaced with a transformer encoder applied directly to tokenized amino-acid sequences, while the remaining training and evaluation protocol was matched to the full model.

### 2.9 Benchmark-Sensitivity Settings

To assess the sensitivity of the evaluation benchmark to negative-set construction and species background, two additional studies were performed alongside the original held-out benchmark. In the first analysis, the proxy-negative selection procedure was replaced with random sampling from a large PeptideAtlas-derived pig peptide pool containing 87,908 unique sequences. This setting isolated the effect of negative-set construction on model performance. Sampled peptides were filtered to satisfy the model-compatible length range, canonical amino acid alphabet, and exclusion of sequences overlapping with the positive and unlabeled sets. In the second analysis, the original proxy negative construction pipeline was retained, but the external candidate pool was changed from *C. elegans* peptides to the same pig peptide background set. This setting evaluated the effect of species background on benchmark performance. In all cases, final performance was evaluated using the ensemble of all six models on the corresponding held-out benchmark, with a fixed threshold selected from validation data only.

## 3 Results

### 3.1 Performance of Pep-PU-GAN

On the held-out benchmark of 810 positive peptides and 810 proxy negatives, the mean prediction of the six-model ensemble achieved a precision of 0.93 (95% CI: 0.909–0.943), recall of 0.93 (95% CI: 0.910–0.946), F1 score of 0.93 (95% CI: 0.914–0.940), AUROC of 0.98, and accuracy of 0.93 (95% CI: 0.914–0.940) at the validation-derived threshold of 0.4951. Table 1 reports model-selection benchmark performance for the six trained models.

**Table 1.**
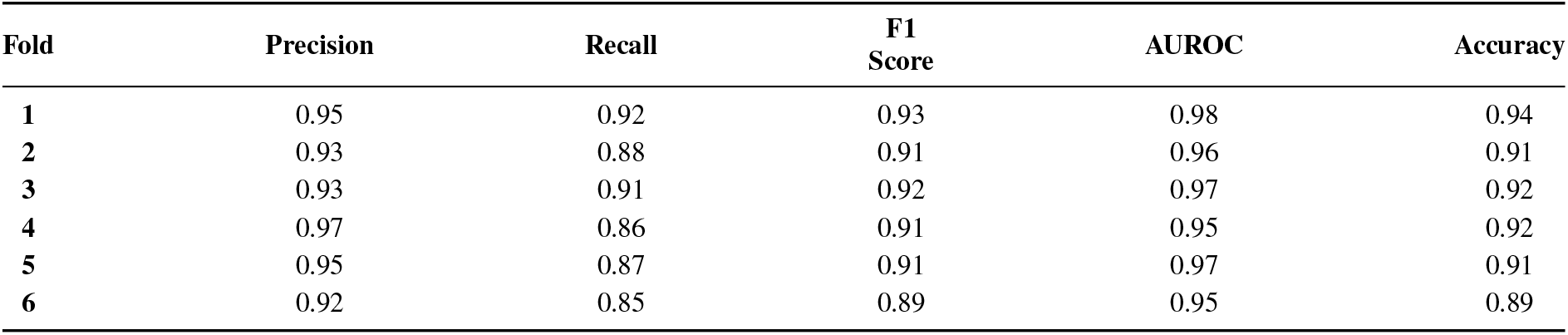
Validation-benchmark performance of Pep-PU-GAN across Folds 1–6.

| Fold | Precision | Recall | F1 Score | AUROC | Accuracy |
| --- | --- | --- | --- | --- | --- |
| 1 | 0.95 | 0.92 | 0.93 | 0.98 | 0.94 |
| 2 | 0.93 | 0.88 | 0.91 | 0.96 | 0.91 |
| 3 | 0.93 | 0.91 | 0.92 | 0.97 | 0.92 |
| 4 | 0.97 | 0.86 | 0.91 | 0.95 | 0.92 |
| 5 | 0.95 | 0.87 | 0.91 | 0.97 | 0.91 |
| 6 | 0.92 | 0.85 | 0.89 | 0.95 | 0.89 |

### 3.2 Training Dynamics Across Curriculum Stages

To evaluate the curriculum learning strategy, we report F1 and AUROC at three stages: epoch 8 (end of PU warm-up), epoch 23 (completion of GAN introduction), and the final epoch per fold (early stopping on validation F1), as detailed in Table 2. Fig. 4 shows the discriminator’s epoch-wise nnPU training loss across the six folds, while Fig. 5 shows the Fold 6 PU-branch representations at initialization, epoch 10, and the best validation-F1 checkpoint at epoch 19.

**Table 2.** F1 and AUROC scores for Fold 1 and Fold 6 at selected stages of curriculum learning (Epoch 8, Epoch 23, and the final training epoch).

| Metric | Epoch 8 | Epoch 23 | Final epoch |
| --- | --- | --- | --- |
| **F1 Fold 1** | 0.87 | 0.86 | 0.93 |
| **F1 Fold 6** | 0.85 | 0.88 | 0.88 |
| **AUROC Fold 1** | 0.94 | 0.94 | 0.98 |
| **AUROC Fold 6** | 0.93 | 0.96 | 0.96 |

**Fig. 4.**
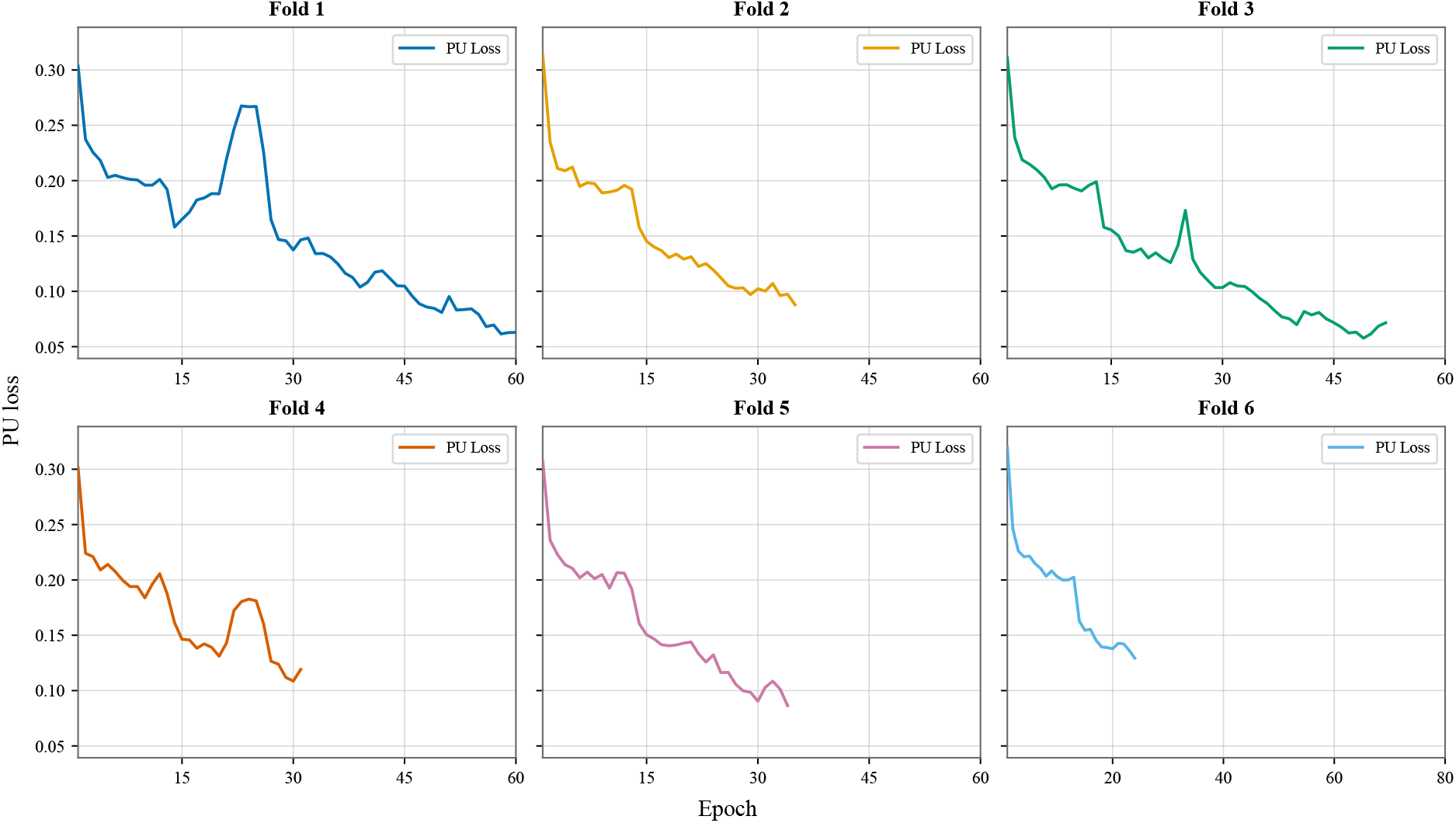
Discriminator nnPU training loss across six folds. Each curve shows the mini-batch-averaged nnPU loss per epoch.

**Fig. 5.**
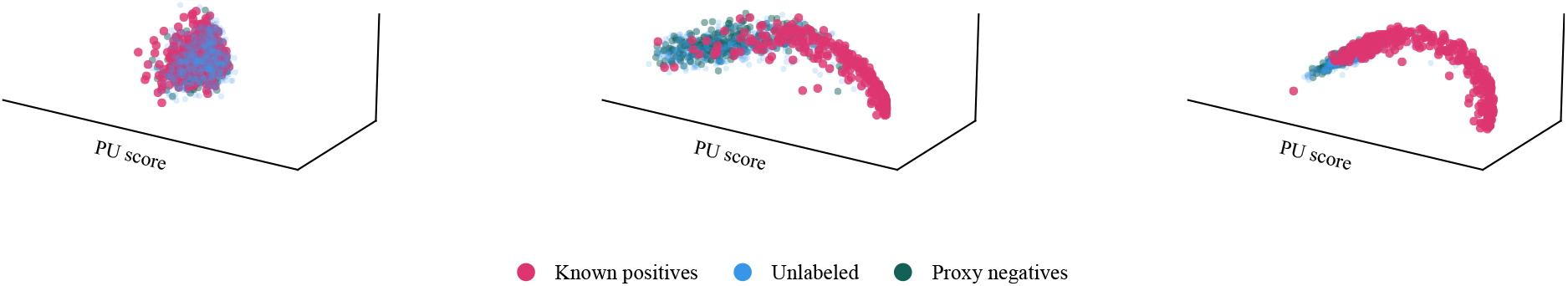
Evolution of the Fold 6 PU-branch representation across training states. The same 324 labeled-positive calibration peptides, 500 peptides sampled from the PU-training unlabeled pool, and 324 calibration proxy negatives are shown in each panel. (A) Model initialization at epoch 0. (B) Intermediate training checkpoint at epoch 10. (C) Best validation-F1 checkpoint at epoch 19. For each panel, the 64-dimensional features produced by the PU branch were standardized, and the horizontal coordinate represents the logit-transformed positive-class probability produced by the PU head. The other two coordinates are the first two principal components of the feature representation after linear removal of the score-associated component. Colors denote labeled-positive calibration peptides, unlabeled peptides, and calibration proxy negatives.

### 3.3 Adaptation of the Positive-Risk Weight

To characterize the adaptation of the positive-risk weight, we report for each fold the initial class-prior estimate 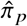, the mean adaptive weight across the executed training epochs, and the final adaptive weight.

As shown in Table 3, the mean adaptive risk weights remained close to their initial values in Folds 2 and 6, whereas Folds 4 and 5 underwent upward adjustments during training before ending below their initial values.

**Table 3.** Fold-wise initial class-prior estimates and adaptive positive-risk weights for Pep-PU-GAN. The mean adaptive weight is computed over all executed epochs within each fold.

| Fold | Initial class-prior estimate | Mean adaptive risk weight | Final adaptive risk weight |
| --- | --- | --- | --- |
| 1 | 0.239 | 0.231 | 0.152 |
| 2 | 0.237 | 0.237 | 0.237 |
| 3 | 0.237 | 0.220 | 0.171 |
| 4 | 0.238 | 0.239 | 0.235 |
| 5 | 0.237 | 0.247 | 0.211 |
| 6 | 0.278 | 0.278 | 0.278 |

### 3.4 Comparison with the GAN-Free Baseline

Comparison with the GAN-free baseline on the independent test set shows that the integrated adversarial and self-training components improved performance:

Table 5 summarizes the initial class-prior estimates and adaptive positive-risk weights across the six folds for the GAN-free baseline.

Together with Table 4, Table 5 shows that the combined GAN and self-training components change the validation signals and, consequently, the fold-wise adaptation trajectories of the positive-risk weight.

**Table 4.**
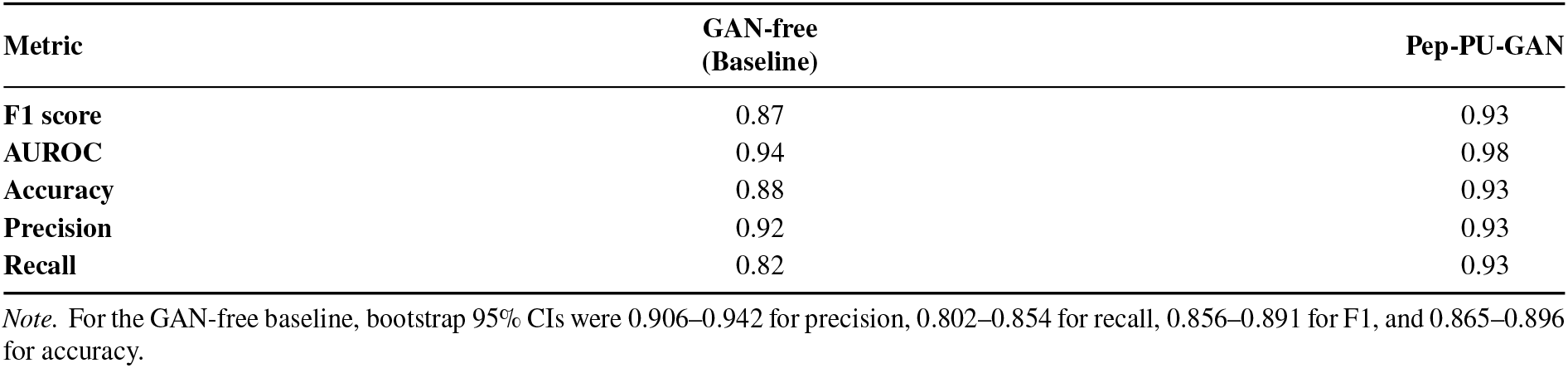
Comparison of GAN-free baseline vs. Pep-PU-GAN on the independent test set.

| Metric | GAN-free (Baseline) | Pep-PU-GAN |
| --- | --- | --- |
| F1 score | 0.87 | 0.93 |
| AUROC | 0.94 | 0.98 |
| Accuracy | 0.88 | 0.93 |
| Precision | 0.92 | 0.93 |
| Recall | 0.82 | 0.93 |

*Note.* For the GAN-free baseline, bootstrap 95% CIs were 0.906–0.942 for precision, 0.802–0.854 for recall, 0.856–0.891 for F1, and 0.865–0.896 for accuracy.

### 3.5 Impact of Peptide Graph Modeling

Removing the sequence-derived graph encoder resulted in a substantial performance decline on the independent test set, with F1 and accuracy decreasing from 0.93 to 0.80 and AUROC from 0.98 to 0.89 (Table 6). This consistent degradation supports the contribution of peptide graph modeling to predictive performance. Fig. 6 further illustrates the differences in accuracy and AUROC among Pep-PU-GAN, the GAN-free PU baseline, and the graph-free sequence baseline.

**Table 5.** Fold-wise initial class-prior estimates and adaptive positive-risk weights for the GAN-free baseline. The mean adaptive weight is computed over all executed epochs within each fold.

| Fold | Initial class-prior estimate | Mean adaptive risk weight | Final adaptive risk weight |
| --- | --- | --- | --- |
| 1 | 0.239 | 0.261 | 0.263 |
| 2 | 0.237 | 0.246 | 0.249 |
| 3 | 0.237 | 0.238 | 0.261 |
| 4 | 0.238 | 0.238 | 0.238 |
| 5 | 0.237 | 0.248 | 0.249 |
| 6 | 0.278 | 0.295 | 0.306 |

**Table 6.** Graph-encoder ablation on the independent test set. Performance metrics compare full Pep-PU-GAN with the graph-free baseline.

| Metric | Full<br>Pep-PU-GAN | Graph-free<br>Ablation |
| --- | --- | --- |
| F1 score | 0.93 | 0.80 |
| AUROC | 0.98 | 0.89 |
| Precision | 0.93 | 0.78 |
| Recall | 0.93 | 0.83 |
| Accuracy | 0.93 | 0.80 |

*Note.* For the graph-free baseline, bootstrap 95% CIs were 0.758–0.802 for precision, 0.810–0.860 for recall, 0.788–0.825 for F1, and 0.780–0.819 for accuracy.

**Fig. 6.**
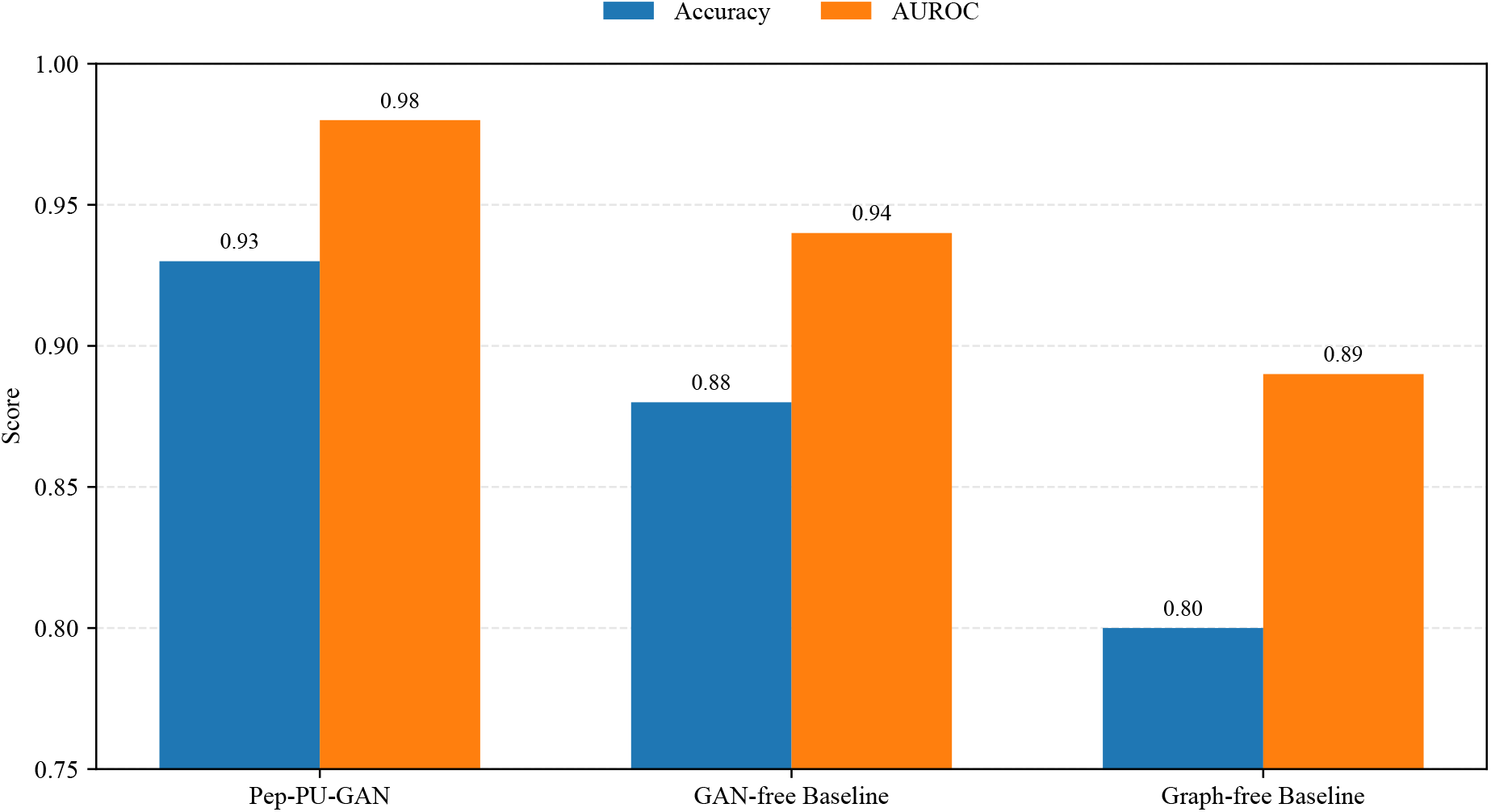
Accuracy and AUROC comparison for Pep-PU-GAN, the GAN-free baseline, and the graph-free baseline.

### 3.6 Sensitivity Analyses of Negative Construction and Species Background

Table 7 summarizes performance across the original benchmark and two sensitivity settings, and Fig. 7 visualizes the corresponding metric changes and positive-risk-weight trajectories. Overall, performance was similar across the three settings, although benchmark composition affected the final metrics. Random pig peptides produced the highest recall (0.96), whereas the original benchmark provided the most balanced performance across metrics. Using pig peptides within the proxy-negative construction pipeline resulted in slightly lower performance, with precision, recall, F1 score, AUROC, and accuracy of 0.90, 0.92, 0.91, 0.97, and 0.91, respectively.

**Table 7.** Comparison of final held-out benchmark performance under the original benchmark and two additional sensitivity analyses.

| Metric | Original<br>(*C. elegans*<br>proxy negatives) | Random pig<br>peptides | Pig proxy<br>negatives |
| --- | --- | --- | --- |
| Precision | 0.93 | 0.89 | 0.90 |
| Recall | 0.93 | 0.96 | 0.92 |
| F1 score | 0.93 | 0.92 | 0.91 |
| AUROC | 0.98 | 0.98 | 0.97 |
| Accuracy | 0.93 | 0.92 | 0.91 |
| Threshold | 0.4951 | 0.5196 | 0.5686 |

Note. Bootstrap 95% CIs (precision, recall, F1, accuracy) were 0.863–0.901, 0.940–0.968, 0.904–0.929, and 0.899–0.927 for random pig peptides, and 0.879–0.917, 0.891–0.930, 0.890–0.919, and 0.889–0.918 for pig proxy negatives.

**Fig. 7.**
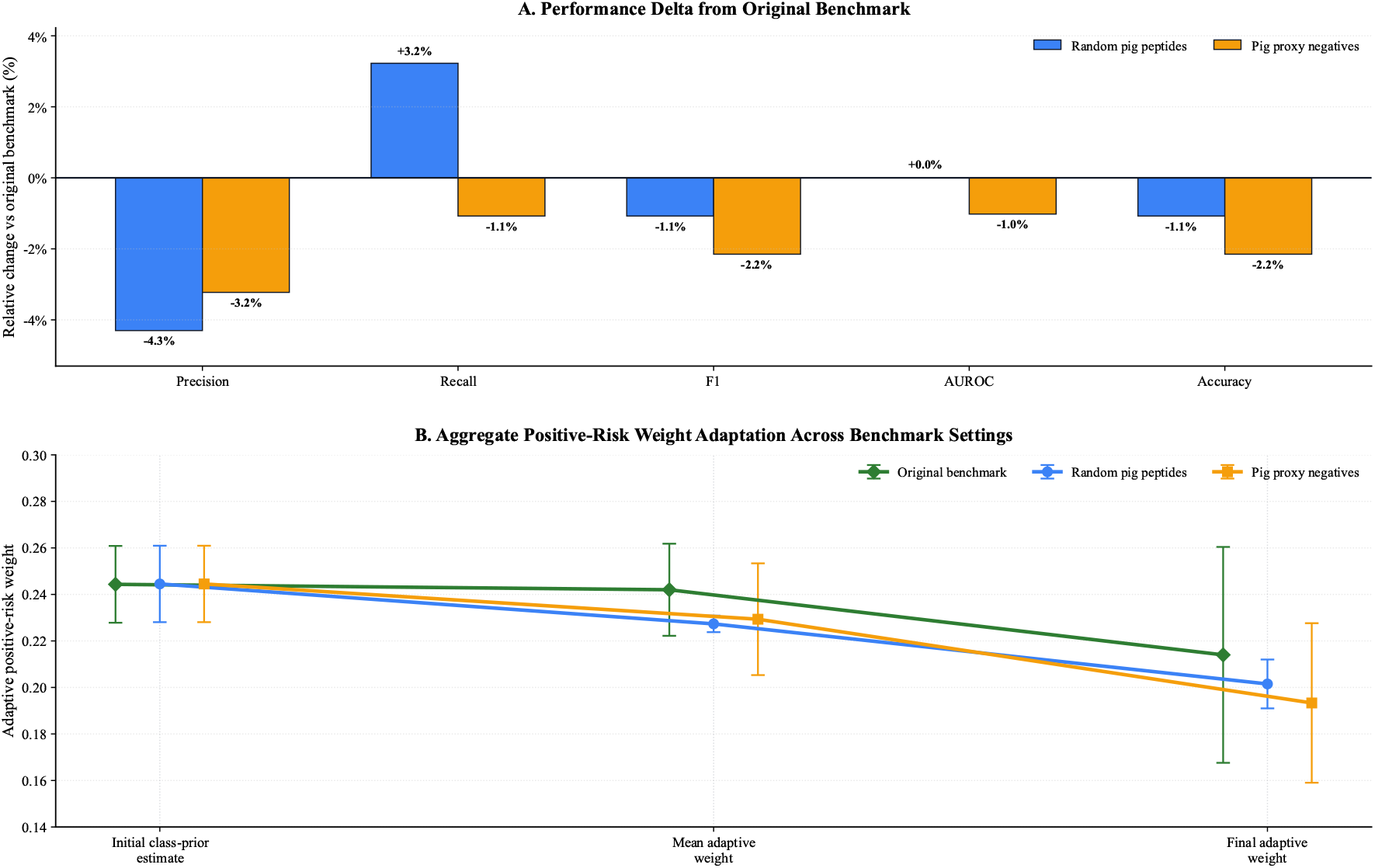
Benchmark sensitivity to negative-set construction and species background. In (A), relative changes in held-out performance are shown for benchmarks constructed with random pig peptides or pig proxy negatives, compared with the original benchmark based on *C. elegans* proxy negatives. Values above and below zero indicate improved and reduced performance relative to the original benchmark, respectively. In (B), validation-guided adaptation of the positive-risk weight is summarized across benchmark settings. Points indicate the mean across six folds for the initial class-prior estimate, mean adaptive risk weight, and final adaptive risk weight; error bars denote the standard deviation across folds.

Table 8 compares the fold-wise initial class-prior estimates and adaptive positive-risk weights for the two additional sensitivity analyses. The initial estimate remained unchanged across both settings because the positive and unlabeled training partitions were identical. In contrast, the mean and final adaptive weights varied because the weight-update rule uses performance on the corresponding validation benchmark. The randomly sampled pig set produced lower final adaptive weights across all folds, whereas the pig proxy-negative setting produced greater fold-to-fold variation.

**Table 8.** Positive-risk weights across benchmark sensitivity analyses. Initial class-prior estimates and adaptive positive-risk weights are shown across six folds. The mean adaptive weight is computed over all executed epochs within each fold.

|  | Fold 1 | Fold 2 | Fold 3 | Fold 4 | Fold 5 | Fold 6 |
| --- | --- | --- | --- | --- | --- | --- |
| *Random pig peptides* |  |  |  |  |  |  |
| Initial class-prior estimate | 0.239 | 0.237 | 0.238 | 0.238 | 0.237 | 0.278 |
| Mean adaptive risk weight | 0.224 | 0.231 | 0.224 | 0.228 | 0.225 | 0.232 |
| Final adaptive risk weight | 0.194 | 0.192 | 0.192 | 0.214 | 0.214 | 0.203 |
| *Pig proxy negatives* |  |  |  |  |  |  |
| Initial class-prior estimate | 0.239 | 0.237 | 0.238 | 0.238 | 0.237 | 0.278 |
| Mean adaptive risk weight | 0.239 | 0.209 | 0.230 | 0.228 | 0.201 | 0.269 |
| Final adaptive risk weight | 0.239 | 0.156 | 0.192 | 0.192 | 0.156 | 0.225 |

### 3.7 Model-Based Characterization of the Unlabeled Peptide Pool

To characterize the distribution of model outputs within the unlabeled peptide pool, we evaluated the PeptideAtlas-derived Cow Peptides set using the trained Pep-PU-GAN ensemble. For each peptide, we calculated the ensemble-mean positive-class score produced by the PU classification branch and the ensemble-mean cosine distance to the nearest labeled positive in the learned embedding space. The positive-class score summarizes the output of the PU classification branch, whereas the nearest-positive cosine distance measures representation-space proximity to the labeled-positive reference set, with smaller distances indicating greater similarity.

As shown in Fig. 8, 1,002 of the 8,558 unlabeled peptides (11.7%) had positive-class scores at or above the predefined decision threshold of 0.4951. Among these 1,002 peptides, 811 had nearest-positive cosine distances below the 75th-percentile distance reference line, whereas 191 had distances at or above this reference line. Thus, the above-threshold peptides were distributed across both lower- and higher-distance regions of the learned representation space. This analysis characterizes how the trained model organizes the unlabeled peptide pool according to its classification scores and learned representations.

**Fig. 8.**
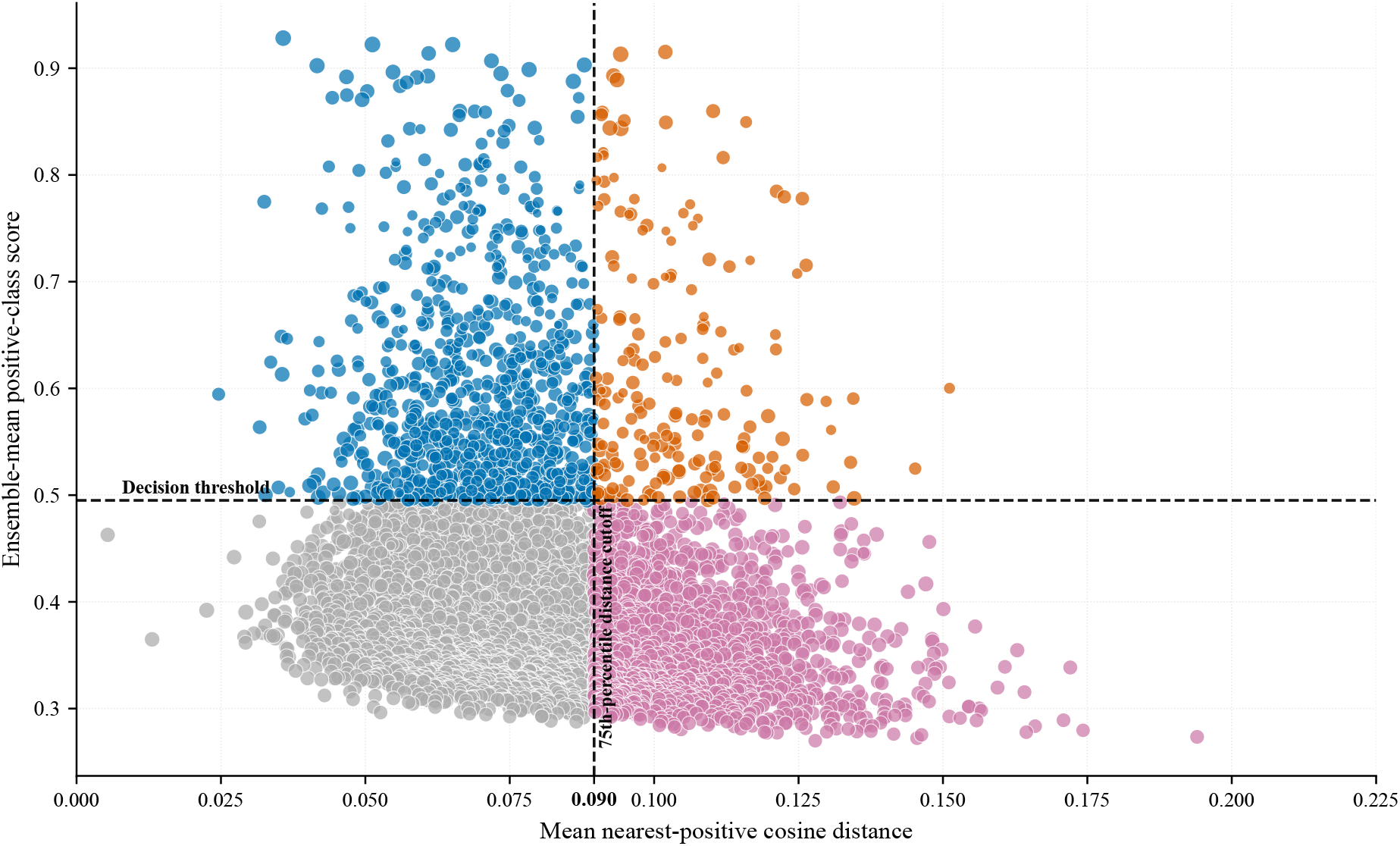
Characterization of the unlabeled peptide pool. The PeptideAtlas-derived Cow Peptides unlabeled pool is characterized using the ensemble-mean positive-class score and ensemble-mean cosine distance to the nearest labeled positive in the learned embedding space. The horizontal dashed line marks the predefined decision threshold of 0.4951; points at or above this line constitute the above-threshold group. The vertical dashed line marks the 75th-percentile reference line of the nearest-positive cosine-distance distribution across the unlabeled pool. Blue and orange points represent above-threshold peptides below and at or above the distance reference line, respectively, whereas gray and magenta points represent below-threshold peptides below and at or above the distance reference line, respectively. Smaller distances indicate greater representation-space similarity to the labeled-positive reference set.

## 4 Discussion

We present and validate Pep-PU-GAN for peptide classification, using neuropeptide identification as a test case where verified negative labels are scarce. Direct comparison with the GAN-free baseline shows clear gains, demonstrating that the full adversarial/self-training framework improves performance over PU training alone. Beyond learning discriminative PU features, adversarial training helps the discriminator detect subtle distributional differences between real and generated embeddings; the hinge loss and feature-matching term appear to stabilize training and improve representation quality. Finally, the self-training loop provides an important practical benefit in refining the decision boundary.

Pep-PU-GAN mitigates GAN instabilities by operating in a continuous latent space, avoiding backpropagation challenges with discrete sequences. Stability is further enhanced by Spectral Normalization in the discriminator’s adversarial branch to constrain the Lipschitz constant (reducing exploding gradients and mode collapse) [26], and generator architectural choices like residual connections for improved convergence and dynamics. Additionally, in our setting, hinge adversarial training with feature matching provided stable optimization in the embedding space.

These results are particularly encouraging because supervised methods typically rely on carefully curated negative examples, which simplifies learning but is often infeasible in peptide domains. These findings indicate that unlabeled peptide data can be informative for peptide identification when combined with PU risk estimation and leakage-controlled benchmark construction.

The graph ablation provides a more specific interpretation of the encoder’s contribution. In this setting, the observed gain reflects local message passing across adjacent residues, attention-based weighting of neighborhood information, and graph-level pooling over context-enriched residue embeddings. Thus, the graph encoder improves local relational modeling of sequence context.

Ansari et al. are among the few groups to apply PU learning to peptide properties. They used a RNN combined with two PU strategies and evaluated four sequence-based tasks (hemolysis, solubility, non-fouling, and SHP-2 binding) on datasets ranging from a few hundred to tens of thousands of sequences, reporting AUROC values up to ≈ 0.93. By contrast, our Pep-PU-GAN attains an ensemble AUROC of *≈* 0.98 on our neuropeptide benchmark. Because the datasets and prediction tasks differ, this comparison is not a strict head-to-head [8].

### 4.1 Contextual Comparison with Supervised Neuropeptide Predictors

Table 9 provides a contextual comparison of Pep-PU-GAN with recently reported neuropeptide predictors. Despite being trained in the more restrictive PU-learning setting, Pep-PU-GAN achieved comparable reported performance.

**Table 9.**
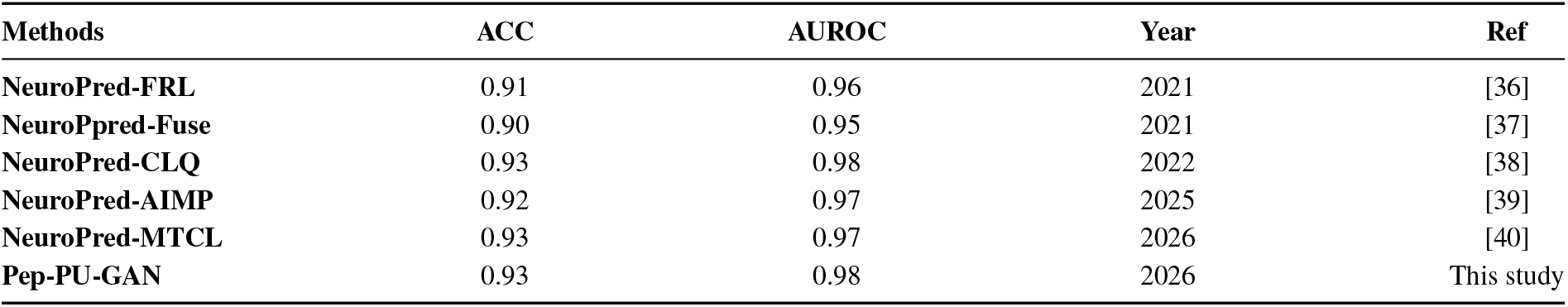
Comparison of Pep-PU-GAN with reported supervised neuropeptide predictors on independent test sets.

### 4.2 Limitations and Future Work

Pep-PU-GAN effectively addresses some PU challenges via GANs but faces limitations: high computational demands for large unlabeled datasets, hyperparameter sensitivity, limited interpretability of learned features, and reliance on distributional assumptions that may not hold in heterogeneous biological datasets. Sample generation quality and diversity also need enhancement.

An additional limitation lies in benchmark construction. The evaluation negatives were proxy negatives selected from an external pool after overlap removal and conservative filtering. Although this is a leakage-safe design, it cannot fully exclude the presence of undiscovered positives.

Future work should prioritize scalability, interpretability, and strategies for mislabeled/corrupted data, including richer graph constructions that incorporate physicochemical or structural relations beyond sequential adjacency, multi-class PU extensions, and active learning integration into self-training for more adaptive, data-efficient refinement.

## 5 Conclusions

This study introduced Pep-PU-GAN, a PU deep-learning framework for peptide classification that combines a sequence-derived residue graph encoder with latent-space adversarial augmentation and generator-assisted self-training. It outperformed the evaluated baselines and achieved performance comparable to that of reported supervised methods on the neuropeptide task. These results demonstrate that Pep-PU-GAN provides an effective framework for peptide-function prediction when verified negative labels are unavailable.

## Declarations

### Availability of data and materials

The Pep-PU-GAN implementation and the data used in this study are publicly available in the GitHub repository:

https://github.com/Farzad-Midjani/Pep-PU-GAN.

The datasets included in the repository were derived from the publicly available peptide sequence resources cited in the manuscript. No proprietary or restricted-access data were used.

### Ethics approval and consent to participate

Not applicable.

### Consent for publication

Not applicable.

### Authors’ contributions

F.M. conceived the study, developed the methodology, curated the data, implemented the computational framework, performed the experiments and formal analyses, interpreted the results, prepared the figures and visualizations, and wrote the original manuscript.

S.H. contributed to methodology, data curation, formal analysis, interpretation of the results, and writing and revision of the manuscript.

F.Z.K. contributed to formal analysis, validation of the computational results, visualization, and review and editing of the manuscript.

M.M. contributed to data curation, validation, and writing and revision of the manuscript.

B.S.A. contributed to validation, interpretation of the results, and review and editing of the manuscript.

B.K. contributed to conceptualization, methodology, interpretation of the results, supervision, project administration, and review and editing of the manuscript, and served as the corresponding author.

All authors reviewed and approved the final manuscript.

## Acknowledgements

Not applicable.

